# Chromosome-scale genome assembly of *Cycas revoluta* provides insights into cycad diversification and species diversity

**DOI:** 10.64898/2026.07.11.737888

**Authors:** Mitsuhiko P. Sato, Yuta B. Aoyagi, Kazutoshi Yoshitake, Yukiho Toyama, Hideaki Iimura, Fumika Matsuoka, Atsushi Toyoda, Takashi Akagi, Satohiro Okuda, Yasuko Ito-Inaba, Kenta Shirasawa

**Author notes:** Department of Frontier Research, National Institute of Genetics, Shizuoka 411-8540, Japan (Yukiho Toyama).

## Abstract

Cycads are an ancient group of seed plants. Despite their ancient origin, many extant cycad genera exhibit high species diversity. The specialized reproductive traits in cycads, dioecy governed by the XY sex-determination system and the elaborate co-evolutionary synergy with insect pollinators, may facilitate lineage diversification. Here, we present a chromosome-scale genome sequence of *Cycas revoluta*, the species in which plant spermatozoids were discovered in 1896. The genome sequence spanned 11.6 Gb, 98.4% of which were anchored onto the 11 cycad chromosomes. Repetitive sequences occupied 9.8 Gb, and 31,481 genes were predicted. Based on this genome assembly, the X- and Y-associated genomic regions were characterized, and candidate genes for sex determination were identified. In addition, the genomic positions of genes for sex-related traits were determined. Subsequently, we analyzed transcriptomes for thermogenesis responsible for attracting insect pollinators. Through these comprehensive analyses, we provide new insights into the genomic basis of cycad diversification and species diversity.

## Introduction

Cycad (order Cycadales) is an ancient group of seed plants with a lineage dating back to the Carboniferous period, approximately 330 million years ago, on the Laurasian landmass and colonized Gondwana in the Jurassic period^1^. Along with cycad, ginkgo (Ginkgoales, a sister order of Cycadales) is also one of the most ancient lineages of extant seed plants. Both lineages diverged from a common ancestor hundreds of millions of years ago. Due to their remarkable morphological resemblance to fossil ancestors, cycad and ginkgo are often termed ‘living fossils.’ Despite their ancient origin, while Ginkgoales comprises only a single extant species, *Ginkgo biloba*, many extant cycad genera exhibit high species diversity resulting from rapid radiations during the Neogene (∼10–20 Ma)^1–4^. The order Cycadales possesses 380 currently known living species, including 125 species in the genus *Cycas*^5^. This raises a central paradox; the coexistence of the long-term genomic stability of an ancient lineage and a species diversification system driven by genomic plasticity in cycad.

A key to understanding this paradox lies in their unique reproductive strategies. The Cycadales and Ginkgoales lineages are characterized by a fertilization system possessing primitive motile sperm like basal plants and derived pollen tubes like seed plants^6^, which intermediate state between zooidogamy and siphonogamy in cycad is supported by molecular evidence through tissue-specific transcriptome analysis^7^. Furthermore, both lineages exhibit strict dioecy governed by stable XY sex-determination systems^8^. Recently, a candidate sex determination gene was identified in the male-specific region (MSR) of the Zamiaceae and *Ginkgo* genomes^2^, with homologous genes found in *Cycas panzhihuaensis*. Elucidating the genomic basis of these traits is crucial for revealing the evolutionary history of reproduction in land plants.

In contrast to the wind-pollinated and species-poor ginkgo, the rapid radiation of cycads was likely driven by an elaborate co-evolutionary synergy with insect pollinators. Cone thermogenesis, a widespread phenomenon in cycads, is closely associated with reproductive function, such as attracting insect pollinators through the release of volatile emissions and/or infrared radiation^9,10^. This physiological trait suggests that while cycads are morphologically conservative, they possess highly specialized physiological adaptations to synchronize with specific biotic partners. Understanding how these specialized reproductive traits facilitated lineage diversification is vital to explaining why Cycadales, unlike their sister order, successfully exploited modern ecological niches through intricate biotic interactions.

To address these questions, a chromosome-scale genome assembly serves as a foundation to reveal chromosome structures and transposon evolution, determine sex-associated regions, and accurately identify gene models and their expression profiles. Owing to the advancement of sequencing technologies and sophisticated bioinformatics tools, high-quality chromosome-scale genome assemblies are now available for many plant species^11^, including gymnosperms^2,12,13^. However, the large genome size and repeat-rich sequences of gymnosperm genomes are still challenging in general. Hi-C scaffolding is a reliable method for building chromosome-scale sequences^14^. Alternatively, genetic mapping is also a promising method to anchor sequences to genetic maps, establishing chromosome-scale sequences even in gymnosperms^13^. For genetic mapping, mapping populations consisting of hundreds of samples should be generated by crossing parental lines, which requires time and space and is not realistic in organisms with large body sizes and/or long life cycles. In gymnosperms, megagametophytes have been used for linkage analysis^15^. In our previous study, we used pollen, each possessing a recombinant haploid chromosome derived from the heterozygous diploid genomes of the parent, as a mapping population to demonstrate that single-pollen genotyping technology is useful to establish genetic maps in plants^16^. These techniques would also be helpful to establish a chromosome-scale genome assembly in cycad.

In this study, we focused on *Cycas revoluta*, a species representing the northernmost limit of cycad distribution, native to southern Japan. Beyond its ecological significance, *C. revoluta* has a profound ethnoecological and symbolic history within the culture of the Ryukyu archipelago^17^. Here, we report a chromosome-scale genome assembly of *C. revoluta*, spanning over 11 Gb in length. By integrating this high-quality genomic resource with tissue-specific transcriptomic data, we uncovered the genetic architecture that facilitates both ancestral preservation and modern adaptive innovation. This study provides a foundational tool for resolving the central paradox of cycad evolution and provides new insights into the genomic basis of their unique reproductive and sex-determination systems.

## Materials and methods

### Plant Materials

Six *Cycas revoluta* individuals were used, including one male and one female from each of University of Miyazaki (Miyazaki, Japan), Kazusa DNA Research Institute (Kisarazu, Japan), and the Koishikawa Botanical Garden of the University of Tokyo (Tokyo, Japan). The male individual from Miyazaki was used for *de novo* genome assembly and the other five lines were used for genetic diversity analysis.

### Whole genome sequencing and de novo assembly

Leaves of the male individual from Miyazaki were subjected to high-molecular-weight genomic DNA extraction using a Genomic-tip 500/G (Qiagen, Hilden, Germany). The high-molecular-weight genomic DNA was sheared to 30 kb using Megaruptor2, and long-read sequencing libraries were prepared using the SMRTbell Express Template Prep Kit 2.0 (PacBio, California, United States). The libraries were separated using BluePippin (Sage Science, Massachusetts, United States) to remove short DNA fragments (< 15kb). Sequence data were obtained using the Sequel II system (PacBio) and converted into high-fidelity (HiFi) reads with the SMRT Link pipeline (PacBio). The HiFi reads were assembled into contig sequences using Hifiasm version 0.19.6^18^, with the parameter that removed 20 bases from both ends of the reads. In parallel, genome DNA was extracted from leaves of the other five individuals and sequenced using the HiFi method on the Revio system (PacBio).

An Omni-C library, a type of high-throughput chromosomal conformation capture (Hi-C)^14^, was prepared using an Omni-C kit (Dovetail Genomics, California, United States) with a standard procedure for scaffolding the contigs. The library was sequenced using DNBSEQ-G400 (MGI Tech, Shenzhen, China). The adapter sequences of Omni-C were trimmed with Cutadapt^19^ and PRINSEQ^20^. Low-quality bases were trimmed with fastx_clipper in the FASTX-Toolkit (http://hannonlab.cshl.edu/fastx_toolkit). The cleaned reads were mapped to the contigs using Bowtie2^21^, and the contigs were scaffolded with YaHS^22^ using default parameters. The contact map was constructed with Juicer^23^ and subsequently visualized and manually curated with Juicebox^24^.

Potential contaminated sequences from organelles and foreign organisms were identified and removed using Tiara v1.0.3^25^ and the gxdb_20240308 dataset of FCS-GX v0.5.5^26^. The completeness of the assembly was evaluated using BUSCO v5.5.0 with embryophyta_odb10^27^ and the gene set of gymnosperms^28^, LTR Assembly Index (LAI)^29^, and telomere_finder^30^. The assembly was aligned to *C. panzhihuaensis* using minimap2 v2.28^31^ and visualized with D-genies^32^.

### Chromosome-scale sequence construction based on SNP analysis

For linkage disequilibrium (LD) analysis, SNPs from pollen were obtained from a single-pollen genotyping technology^16^. Pollen collected from the Miyazaki male individual was subjected to protoplast preparation using the previously reported protocol^33^. Briefly, the pollen of *C. revoluta* was washed with pre-buffer (1/2 MS, 1M sorbitol, 0.5M glucose, 5mM 2-morpholinoethanesulphonic acid, pH 5.8, through a 0.22 µm filter) and filtered through a cell strainer with a 70 µm pore size (pluriSelect Life Science, Saxony, Germany) twice. The washed pollen was soaked in the pre-buffer containing 1.0% Macerozyme R1-0 (Cosmo Bio, Tokyo, Japan), 2.0% Cellulose Onozuka RS (Cosmo Bio), 0.1% BSA as enzyme solution and then shaken at 28D and 20 rpm for 6 days. The pollen protoplasts were collected by centrifugation at 100g for 5 min at room temperature and washed with the pre-buffer twice. Single-cell cDNA libraries were prepared using DNBelab C4 (MGI Tech) and sequenced on DNBSEQ-G400 to generate 150 bp paired-end reads. The obtained reads were mapped to the scaffold sequences as a reference through the standard pipeline of DNBelab C Series Single-Cell RNA Analysis Software (https://github.com/MGI-tech-bioinformatics/DNBelab_C_Series_scRNA-analysis-software) and SNPs were detected with GATK HaplotypeCaller^34^. LD analysis was performed with SELDLA^35^, in which pairwise correlations between loci were calculated using Fisher’s exact test.

For genetic mapping analysis, we used megagametophytes, which were collected from the Kisarazu female individual, as a mapping population. Genomic DNA was extracted from the megagametophytes using the hexadecyltrimethylammonium bromide (CTAB) method^36^ and subjected to ddRAD-Seq library construction using the PstI and MspI enzymes^37^. ddRAD-Seq reads were obtained using DNBSEQ-G400, and their low-quality bases and adaptor sequences were trimmed with PRINSEQ and fastx_clipper in the FASTX-Toolkit, respectively. The cleaned reads were mapped to the genome assembly using Bowtie2^21^, and sequence variants were called using BCFtools^38^. High-confidence SNPs were selected using VCFtools^39^ (parameters for minimum depth of 5; minimum SNP quality of 20; and minor allele frequency of 0.5) and subjected to linkage analysis using Lep-Map3^40^. The resulting genetic map was assigned to the genome assembly using ALLMAPS^41^ to validate the accuracy of the genome assembly.

### Repetitive sequence analysis

Repetitive sequences were detected with RepeatMasker v4.1.5 (https://www.repeatmasker.org) using repeat sequences obtained from the genome assembly using RepeatModeler v2.0.5 (https://github.com/Dfam-consortium/RepeatModeler) and from a dataset registered in Repbase42.

### Protein-coding gene prediction and annotation

Total RNA was isolated from a young leaflet, a mature leaflet, a rachis, a central axis, microsporangia, an upper microsporophyll, and a lower microsporophyll of the Miyazaki male individual with RNA extraction kit (Qiagen). A library for short-read sequencing was prepared with a TruSeq RNA Library Prep Kit v2 (Illumina, California, United States) and sequenced using HiSeq 4000 (Illumina). Potential protein-coding genes were predicted using BRAKER3 version 3.0.7^43^ with the RNA-Seq data, protein sequences of Viridiplantae from orthoDB v11^44^, and a gene set of gymnosperms^28^.

In parallel, a long-read sequencing library was prepared using the KINNEX full-length RNA kit (PacBio) from RNA of a leaflet, a rachis, three stages of microsporangia, and three stages of microsporophyll of the Miyazaki male individual. The library was sequenced on the Revio system (PacBio). The obtained full-length cDNA reads were treated with the Iso-Seq pipeline (https://isoseq.how, PacBio), in which the reads were aligned to the genome assembly with an intron size of 500 kb, to obtain transcript sequences. Since multiple isoforms were aligned to the same genomic region, we selected a single representative transcript per locus based on the following criteria: (1) support by more than three reads; (2) the presence of an open reading frame; and (3) a splicing pattern matching either the gene models of *C. panzhihuaensis* or those predicted by the RNA-Seq analysis described above. For genomic regions where no gene models were assigned, the transcript with the highest number of reads was selected.

The genes predicted with short- and long-read sequencing were compared using GffCompare^45^ and single representative genes per locus were selected from the long-read data with highest priority followed by the short-read data. The predicted genes were assigned functional annotations with eggNOG-mapper v2.1.8^46^ and clustered with those of *C. panzhihuaensis*^2^ using OrthoFinder^47^.

### Organelle genome assembly and gene prediction

For the six individuals, chloroplast and mitochondrial genome sequences were assembled with oatk v1.0^48^ and the number of repetitive motifs was manually curated by alignment of HiFi reads using GraphAligner^49^. Using Lifton v1.0.5^50^, protein-coding genes and ribosomal genes in the chloroplast assembly were transferred from *C. revoluta* (NC_020319.1) and those in the mitochondrial assembly were transferred from *C. taitungensis* (NC_010303.1), *Ginkgo biloba* (NC_027976.1), *Welwitschia mirabilis* (NC_029130.1), *Pinus Taeda* (NC_039746.1), *Taxus chinensis* (NC_069591.1), *T. wallichiana* (NC_072607.1), and *Abies koreana* (NC_071216.1). Transfer RNA genes were predicted using tRNAscan-SE2.0^51^ and ARAGORN^52^ in both chloroplast and mitochondrial genomes.

### Sequence and structure variations within the Cycas species and across its relatives

HiFi reads from the six individuals were mapped to the *C. revoluta* genome assembly with pbmm2 v1.17.0. Single-nucleotide variations (SNVs) and structural variants (SVs) were called with BCFtools v1.21^38^ and pbsv v2.11.0 (https://github.com/PacificBiosciences/pbsv), respectively. The effects of the variants on gene function were annotated using SnpEff v4.3^53^. The sequence depth for each sample across the genome was calculated using goleft v0.2.6 (https://github.com/brentp/goleft). Homology and synteny analyses between the genomes of *Cycas* and *Ginkgo* were performed using D-genies^32^ and MCScanX^54^, respectively.

### Phylogenetic analysis of MADS-box genes

To identify MADS-box genes from the gene models, we aligned SRF-TF (PF00319) and K-box (PF01486) from the Pfam database (https://www.ebi.ac.uk/interpro/) against the gene models of *C. revoluta* (this study), *C. panzhihuaensis*^2^, and *G. biloba*^12^ with hmmsearch in HMMER v3.3.2 (http://hmmer.org/). Protein sequences of the resultant selected genes in addition to two Y chromosome-specific MADS-box genes, GbMADS18 (MN548764) and CpMADSY (CYCAS_034085) were aligned with MAFFT v7.505^55^. A phylogenetic analysis was conducted using IQ-TREE v3.0.1^56^ with 10,000 ultrafast bootstrap replicates.

### Comparison of candidate genes involved in sex determination and sexual dimorphisms

Toyama et al.^7^ explored gene expression in pollen tubes and capacitated motile sperm of *C. revoluta* in a tissue-specific transcriptome analysis to reveal the intermediate evolutionary state of motile sperm and pollen tubes. The transcriptome analysis was performed without genome sequence data, which were unavailable at the time of analysis. Now, in this study, since we present the *C. revoluta* genome, we aligned a total of 77,449 transcript sequences for nine tissues to the genome sequence as a reference using BLASTX searches to determine their genome positions.

To identify putative genes involved in the biosynthesis and regulation of flagella, 58 flagellar-related genes of six categories (radial spoke protein, central pair, dynein, inner dynein, outer dynein, and intraflagellar transport protein) were collected from a previous study in algae^57^. Similarity searches of the flagellar gene sequences were performed using DIAMOND^58^ against the predicted genes in the genomes of *C. revoluta* (from this study) in addition to 13 land plant species (*Azolla filiculoides*, *Salvinia cucullata*, *G. biloba*, *C. panzhihuaensis*, *Gnetum montanum*, *Sequoiadendron giganteum*, *Abies alba*, *Picea abies*, *Pinus lambertiana*, *P. taeda*, *Pseudotsuga menziesii*, *Oryza sativa*, and *Arabidopsis thaliana*). We employed the reciprocal best hit method with an E value cut-off of 1 × 10^-6^ to select the top hit genes.

### Transcriptome analysis for thermogenesis

Microsporophylls (ML), thermogenic cone tissue, and microsporangia (MG), non-thermogenic cone tissue, were sampled at early thermogenic, middle thermogenic, and late non-thermogenic stages during the flowering period in 2024 from three flowers of the individual used for genome sequencing. The thermogenic tissues and timing in these samples were identified using FLIR thermal image cameras. Total RNA was extracted with an RNA extraction kit (Qiagen), prepared for the library using TruSeq Stranded mRNA HT Sample Prep Kit, and sequenced with DNBSEQ-G400. The resulting reads were mapped to the genome assembly and quantified using hisat2 v2.2.1^59^, stringtie v2.2.1^60^, and edgeR v4.6.3^61^. Mitochondrial transit peptides were predicted from aligned transcripts using TargetP^62^.

## Results

### Chromosome-scale genome assembly

The genome sequence of *C. revoluta* was *de novo* assembled using 17 million HiFi reads totaling 397 Gb (∼29.6x coverage of the estimated genome size of 13.4 Gb^63^), which were obtained from 14 SMRT cells on the Sequel II platform. The reads were assembled into 2,483 contigs spanning 11.6 Gb, with a contig N50 of 14.1 Mb (Table 1). Then, we generated 2.8 billion Omni-C short reads (426 Gb; 31.8x coverage) to scaffold the contigs. Following the mapping of 74.0% of these reads, contigs were scaffolded into 821 sequences. Then, 408 potential contaminated sequences from organelle and foreign organisms were removed. As a result, nine large scaffold sequences exceeding 100 Mb were obtained. Since *C. revoluta* has 11 basic chromosomes, the scaffold sequences might include over-scaffolding or misassemblies.

**Table 1.** Statistics of the *C. revoluta* genome assembly.

|  | Contig sequence | Genome sequence<br>(CREV_r1.0.genome) | Chromosome-scale<br>sequence<br>(CREV_r1.0.pmol) |
| --- | --- | --- | --- |
| Total size (bp) | 11,598,888,748 | 11,582,440,925 | 11,413,479,678 |
| No. of sequences | 2,483 | 334 | 11 |
| N50 length (bp) | 14,138,399 | 1,098,390,435 | 1,098,390,435 |
| Longest sequence size<br>(bp) | 176,690,324 | 1,348,290,124 | 1,348,290,124 |
| No. of gap | 0 | 1,756 | 1,691 |
| Gap length (bp) | 0 | 175,600 | 169,100 |
| No. of predicted genes | Not analyzed | 31,481 | 31,084 |

To address these potential errors, we employed a LD approach. Since pollen grains possess haploid genomes containing recombinant parental chromosomes, haplotype blocks could be predicted using SNPs derived from pollen. We obtained transcriptome sequence data from 210 individual pollen grains and mapped the reads to the scaffold assembly to detect 283 SNPs (Supplementary Figure S1). LD analysis of these SNPs indicated distinct misassembly points in the two largest sequences, which were subsequently broken at these high-confidence junctions.

The accuracy of the resulting sequences was further validated using a genetic mapping strategy. We genotyped 880 SNPs across 224 megagametophytes to construct a genetic map. The resulting map comprised 12 linkage groups with a total length of 1,457.6 cM (Supplementary Figure S2). Ten of the linkage groups were each assigned to a single scaffold sequence, while the remaining two linkage groups were assigned to a single large scaffold, confirming its continuity.

This integrated approach successfully established 11 pseudomolecule sequences at the chromosome-scale (named CREV_r1.0.pmol: Figure 1) together with 323 unplaced contigs, representing the haploid genome of *C. revoluta* (CREV_r1.0.genome: Table 1). The complete BUSCO score was 97.1%, with single-copy and duplicated scores of 91.3% and 5.8%, respectively. The whole-genome LAI score was 15.15. Telomeric repeats (TTTAGGG) were identified at the 15 ends of the 11 chromosome-scale sequences, out of which 5 motifs at one end of 5 chromosome sequences included both the repeats of TTTAGGG and its variant type, TTCAGGG (Supplementary Figure S3).

**Figure 1.**
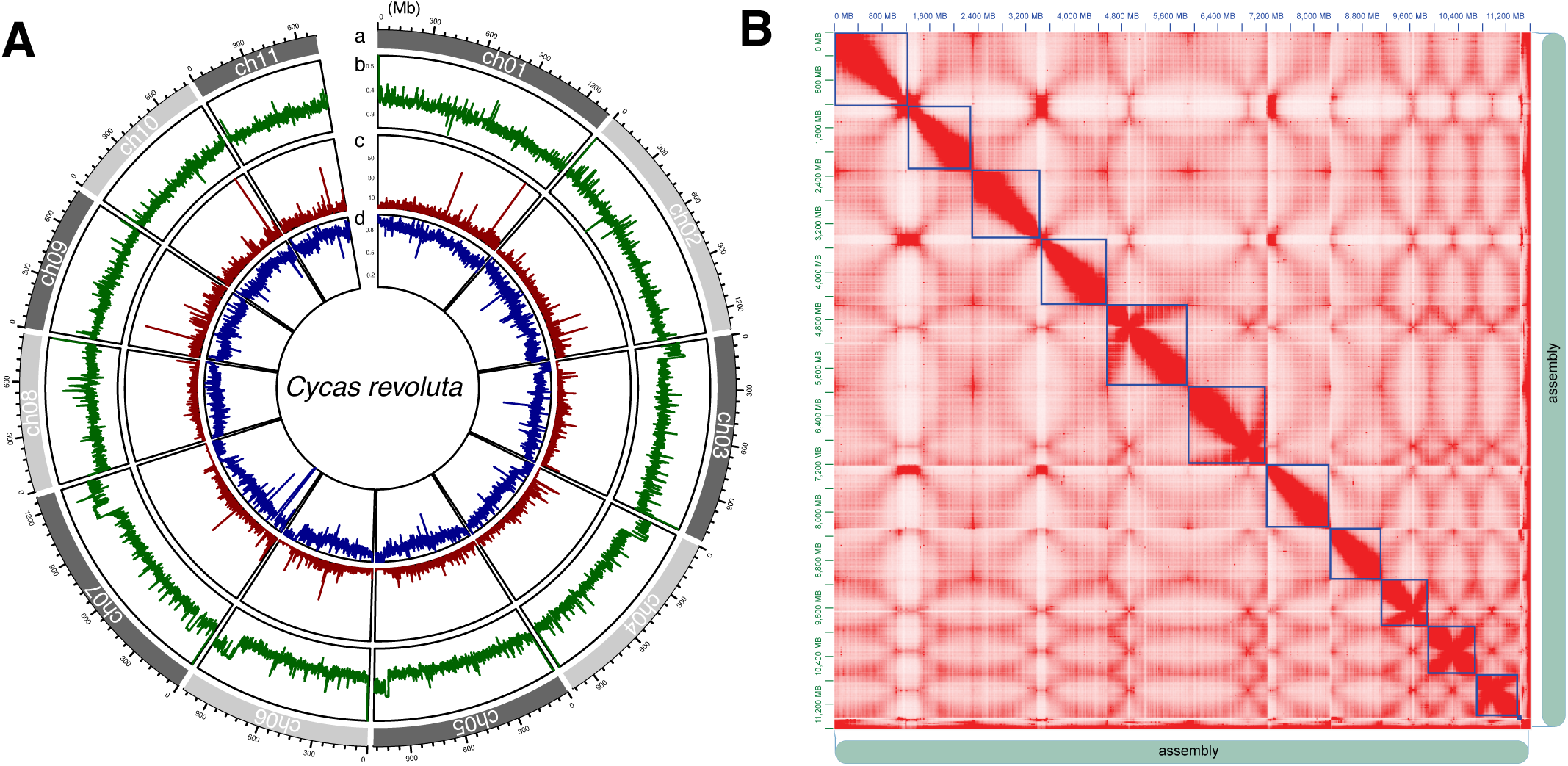
A chromosome-scale genome assembly of *C. revoluta*. (A) A Circos plot of the pseudomolecule assemblies with 1 Mb bins. The tracks from the outside to the inside are (a) pseudo-chromosome numbers, (b) GC content, (c) the number of genes, and (d) repeat density. (B) Omni-C contact map. The color indicates the intensity of interaction, ranging from low (white) to high (red). Blue boxes indicate pseudomolecule assemblies.

The structure of the *C. revoluta* genome was consistent with that of *C. panzhihuaensis*^2^ (Figure 2A). The orientations of the chromosomes are described based on the genomic coordinates, with the 5’ end corresponding to the start of the assembled sequence and the 3’ end to its terminus.

**Figure 2.**
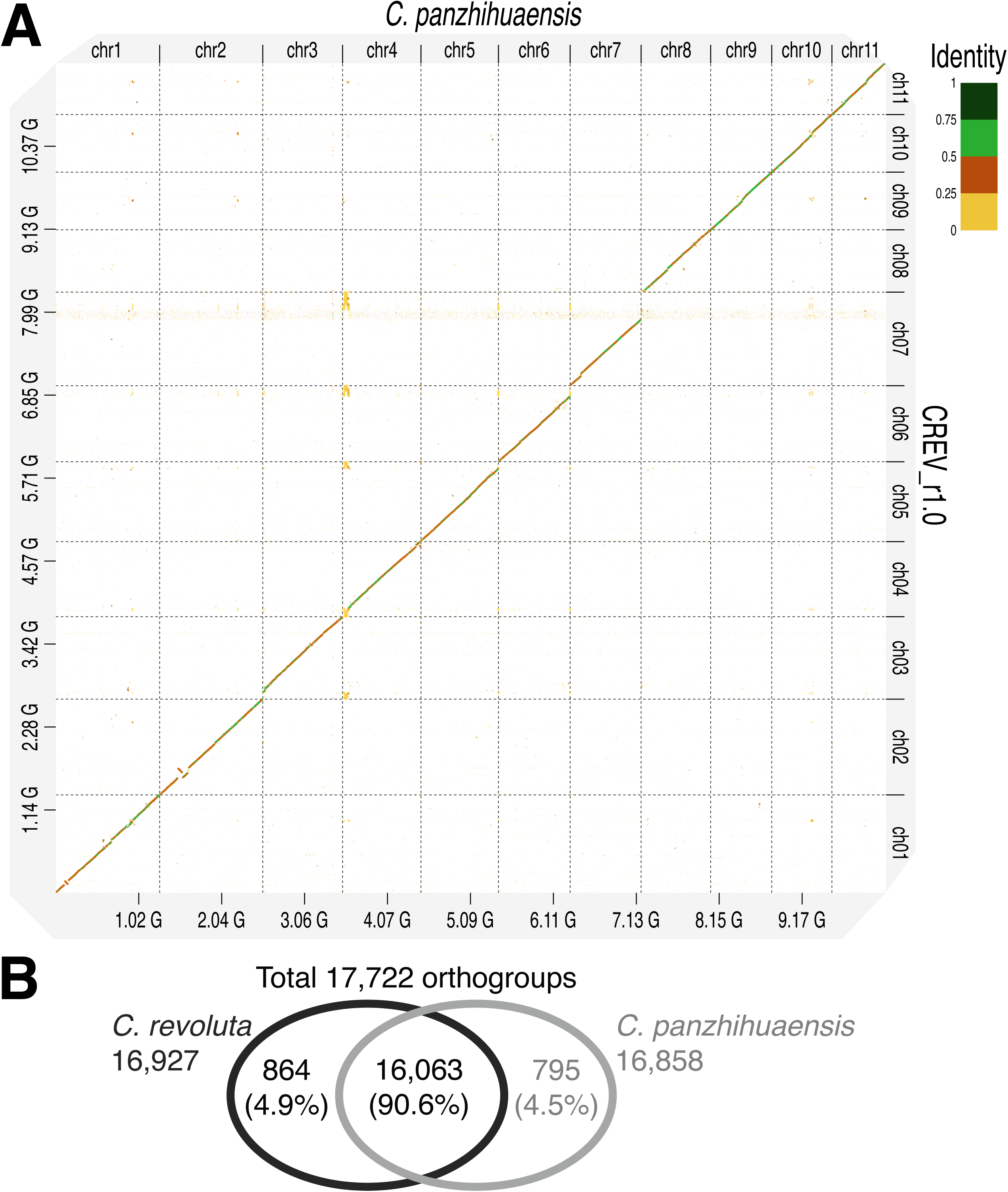
Comparison between *C. revoluta* and *C. panzhihuaensis*. (A) A dot plot between the pseudomolecules of the two species. Line colors represent sequence identity. (B) A Venn diagram of gene clusters.

### Repetitive sequence analysis

Repetitive sequences occupied 9.82 Gb (84.62%) of the 11.60 Gb assembly (Supplementary Table S1). The dominant repeat types were long terminal repeat (LTR) retroelements, which accounted for 51.85% of the total genome, followed by unclassified repeats (20.42%). In contrast, DNA transposons represented only a small fraction of the genome at 3.86% (447.9 Mb).

### Gene prediction

Full-length cDNA sequences were obtained from eight tissues using KINNEX technology. Subsequent genome-guided analysis, which involved mapping these reads and collapsing redundant sequences, yielded 66,769 transcripts. In parallel, RNA-Seq data obtained from seven samples were mapped to the genome sequence. Gene prediction was performed by integrating the non-redundant full-length cDNA sequences with 21,221 genes predicted from RNA-Seq and protein evidence. A total of 31,481 representative protein-coding sequences were identified and functionally annotated with a complete BUSCO score of 87.5%.

Gene clustering analysis between *C. revoluta* and *C. panzhihuaensis* yielded 17,722 orthogroups, of which 16,063 were shared between both species, 864 were unique to *C. revoluta*, and 795 were unique to *C. panzhihuaensis* (Figure 2B). The number of genes between the two species was concordant in 12,340 orthogroups (69.63%), while species-specific gene family expansions or contractions were observed in 137 for *C. revoluta* and 140 for *C. panzhihuaensis*.

### Organelle genome assemblies and polymorphism analysis

The organelle genomes of the six individuals were assembled as single circular sequences in both chloroplasts and mitochondria. The chloroplast genome of the Miyazaki male individual was 162,174 bp in length and encoded 87 protein-coding sequences, 39 tRNA, and 8 rRNA genes. Among the six individuals, a single base insertion/deletion was found in homopolymer stretches, resulting in a total length difference of 1 bp. The mitochondrial genome of the Miyazaki male individual was 417,531 bp in length and encoded 41 protein-coding sequences, 33 tRNA, and 3 rRNA genes. Among the six individuals, nine indels were found, of which six were single base indels whereas the others were variants in the number of tandem repeats in the intergenic region, ranging from 26 to 40 repeats of a 63-bp unit.

### Genetic variations within the species

Genome sequence and structural variations were assessed using the HiFi reads from three male and three female individuals. We identified 10.57 million SNPs and 145,400 structural variants (SVs) over the genome. Both SNPs and SVs were significantly enriched in intergenic regions compared to genic regions (chi-square test, *P*-value < 1.0 x 10^-16^). Even in genic regions, insertions and deletions of 5–20 kb in size were predominantly associated with LTR elements (Figure 3A), suggesting that recent retrotransposon activity continues to shape the genetic landscape of *C. revoluta*. The allele frequency distribution indicated a high prevalence of minor alleles at many loci, reflecting a diverse genetic reservoir within the species (Figure 3B).

**Figure 3.**
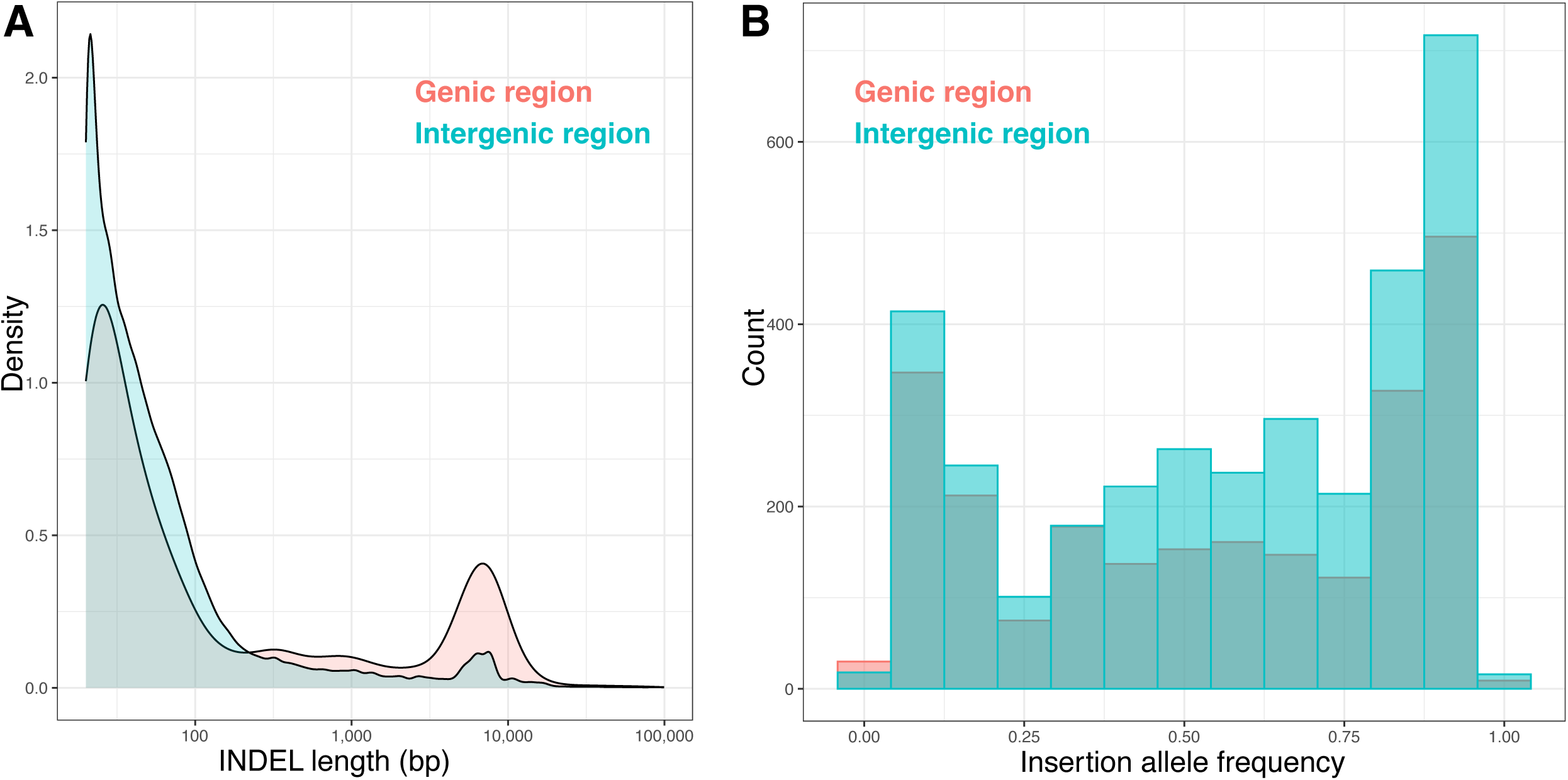
Structural variants within *C. revoluta*. (A) Distribution of INDEL length. (B) Insertion allele frequency of large INDELs (5,000 - 20,000 bp).

### Comparative analysis of genome structures across the relatives

Comparative genomic analysis based on gene synteny revealed a highly conserved chromosome structure between *C. revoluta* and *C. panzhihuaensis* (Figure 4). This macrosynteny indicates that the large-scale genomic architecture is fundamentally maintained within the genus *Cycas*. Based on this structural conservation, the sex-determining region in *C. revoluta* was predicted to be located at the 5’ end of chromosome 8, corresponding to the locus previously identified in *C. panzhihuaensis*. Furthermore, despite their deep phylogenetic divergence, large-scale synteny blocks were also observed between *C. revoluta* and *G. biloba* (Figure 4).

**Figure 4.**
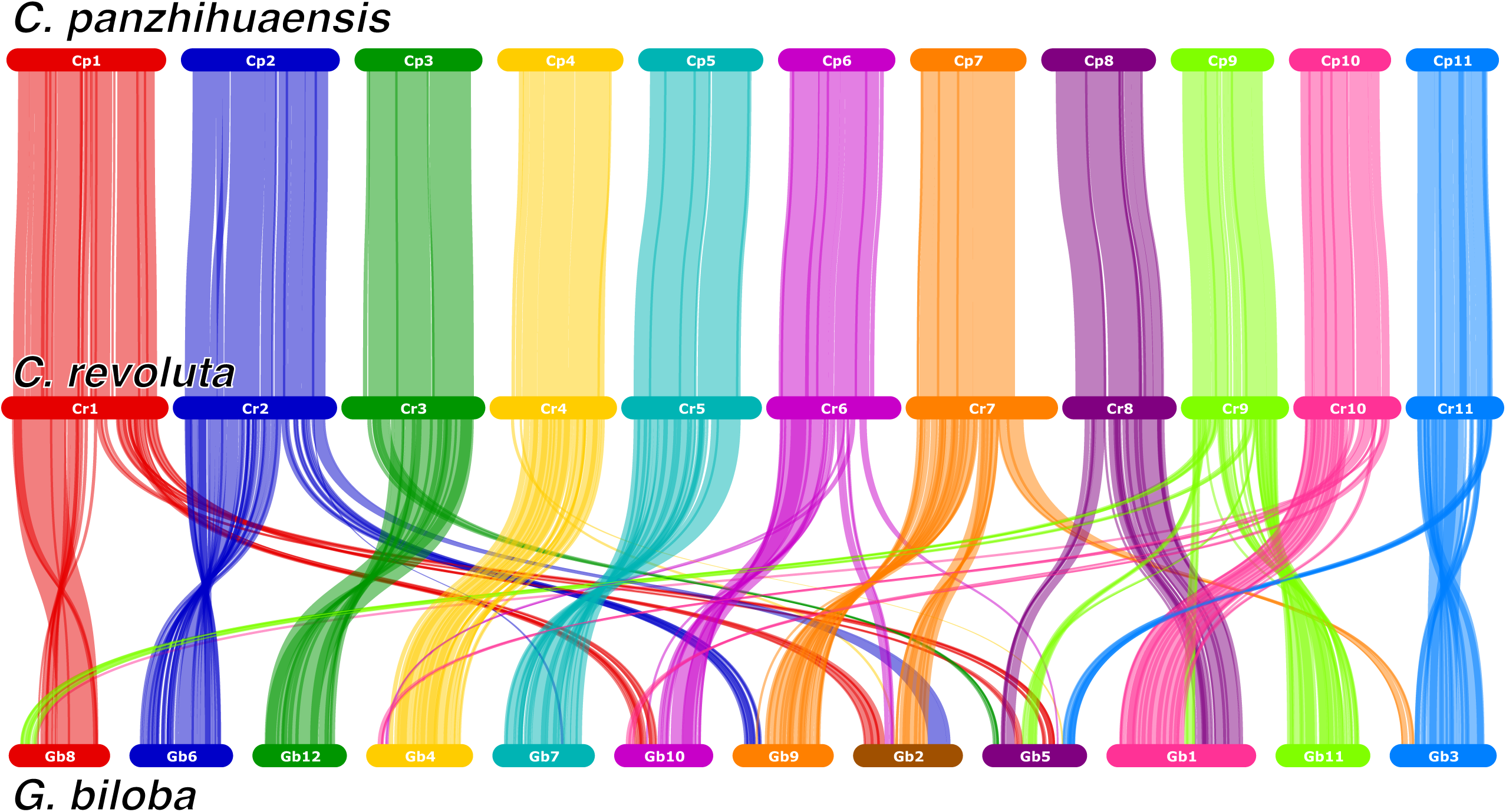
Synteny and collinearity of genes in *C. revoluta*, *C. panzhihuaensis*, and *G. biloba*.

### Identification of sex-associated genomic regions and genes

Although we assembled the genome of a male individual (XY), the assembled sequences would represent either the X- or Y-associated region only. To identify and characterize the sex-specific region on the *C. revoluta* chromosome 8, we performed a depth-of-coverage analysis using the HiFi reads from the male and female individuals (Figure 5). In the three male samples, the depth-of-coverage on the 5’ end of chromosome 8 was half compared to other regions which represent pseudoautosomal regions (PARs) or autosomes (Supplementary Figure S4). On the other hand, in the female samples, the 5’ end of chromosome 8 was rarely mapped with reads while pseudoautosomal regions (PARs) or autosomes were uniformly covered (Supplementary Figure S4). This result suggested that the sequence of the 5’ end of chromosome 8 is the Y-associated region, also known as the male-specific region (MSR) and that sequences for the X-associated region might be missing from the pseudomolecule sequences.

**Figure 5.**
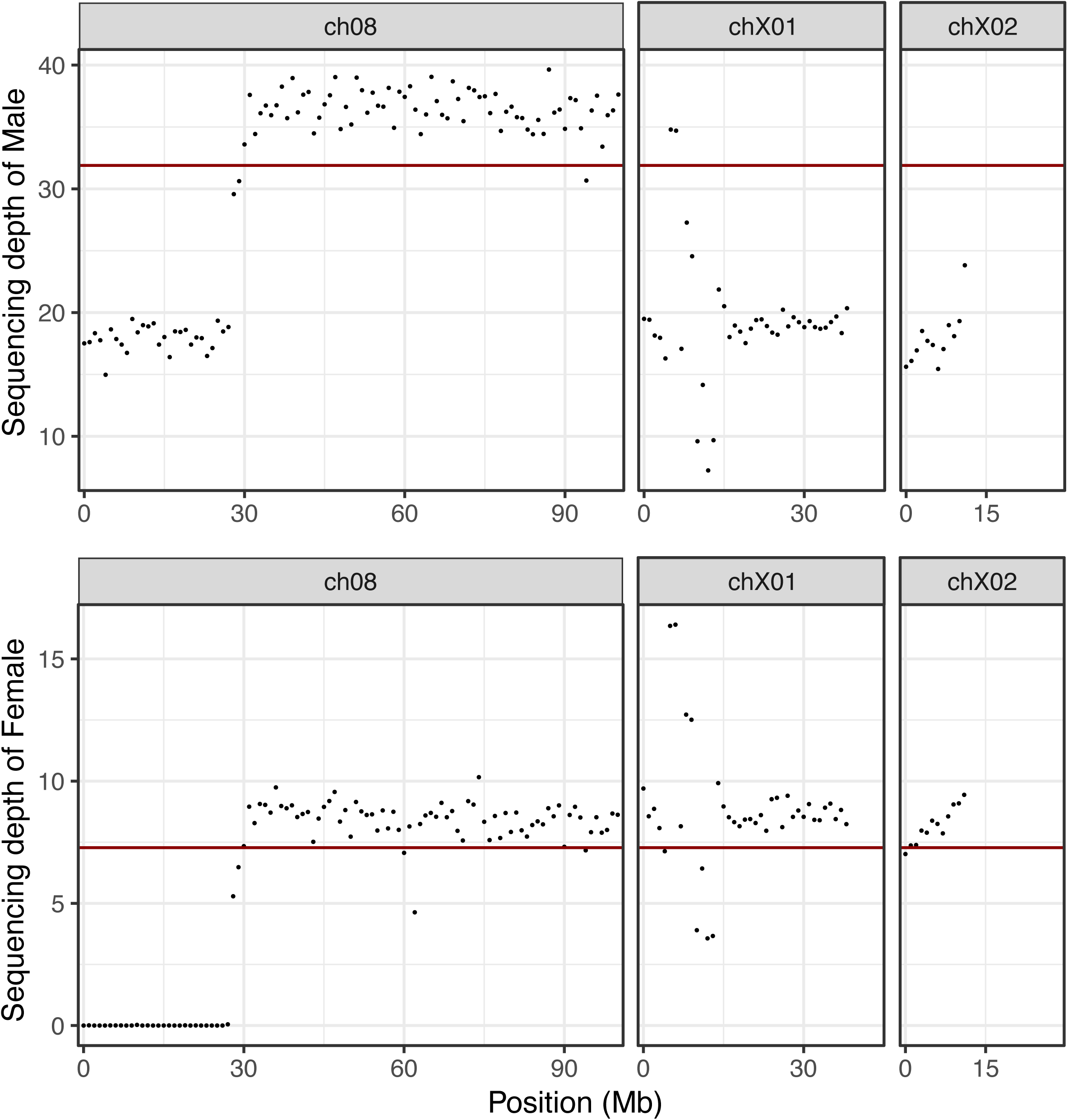
Sequencing depth on sex-determining regions of male and female individuals from Miyazaki. Red lines indicate the mean depth of the whole genome.

In parallel, genome sequences for the X-linked region were recovered from alternate contigs that were derived from alternative haplotypes of the *C. revoluta* diploid genome. First, the alternate contigs were aligned to the genome sequence of *C. panzhihuaensis*, which includes a 124-Mb X-associated region at the end of chromosome 8. As result, two contigs showed high homology to the *C. panzhihuaensis* X- associated region but low homology to the *C. revoluta* Y-linked region (Supplementary Figure S5). Then, the depth-of-coverage analysis was applied to these contigs. As expected, the depth-of-coverage on these contigs was half in the three male samples compared to other regions whereas it was even across these contigs and other regions in the three female samples (Figure 5, Supplementary Figure S4).

Subsequently, we searched for characterized genes within the sex-associated regions. We identified a gene encoding a MADS-box transcription factor (Cre1.0ch08.g000010.1) as a prime candidate for sex determination. This gene shares high sequence similarity with known potential sex-determining genes in basal gymnosperms, specifically CYCAS_034085 (MADS-Y) in *C. panzhihuaensis* and GbMADS4 in *G. biloba*. A genome-wide survey identified 32, 28, and 47 MADS-box genes in *C. revoluta*, *C. panzhihuaensis*, and *G. biloba*, respectively (Supplementary Figure S6). Additionally, a *HOTHEAD*-like gene (Cre1.0ch08.g000080.1) was located within the sex-determining region; its orthologs in *Arabidopsis thaliana* are well-known to cause abnormal floral organ development when mutated^64^. Although the expression of neither the MADS-box nor the *HOTHEAD*-like gene was captured in the above male and female organ transcriptome datasets, their conserved functions and evolutionary positioning strongly support them as key candidates governing sex determination in *C. revoluta*.

### Genes for sex-related traits

To identify genes associated with sex-related traits, we mapped 19,295 unique publicly available transcriptome sequences derived from six tissues^7^ onto the *C. revoluta* genome assembly. Among these, the genomic positions of 16,668 transcripts were successfully determined. Notably, five transcripts were located within the Y-associated male-specific region (MSR), while only one was mapped to the X-associated sequences. Within the Y-associated MSR, three transcripts (Cre978_c2_g1_i1.p1, Cre103_c0_g3_i2.p1, and Cre39109_c0_g1_i2.p1) were exclusively transcribed in male organs, making them strong tissue-specific candidates for sex determination. The remaining two Y-associated transcripts, Cre6872_c0_g1_i1.p1 and Cre94321_c1_g1_i1.p1, were expressed in female organs and megasporophylls, respectively. Conversely, the single X-associated transcript (Cre160992_c0_g1_i1.p1) was expressed in male organs. Among the three male-specific Y-associated transcripts, Cre103_c0_g3_i2.p1 overlapped with a predicted gene model (Cre1.0ch08.g000220.1) encoding a small nuclear ribonucleoprotein, whereas the other two lacked any overlapping gene models or functional annotations.

In parallel, we searched for genes involved in flagellar formation across 14 vascular plant species including 2 angiosperms, 10 gymnosperms, and 2 ferns. The genes for flagellar biosynthesis and regulation, such as radial spoke, central pair, intraflagellar transport, and inner and outer dynein, were absent from the Y-associated region of chromosome 8. Over the genome, the copy numbers of the genes were higher in *Cycas*, *Ginkgo*, and ferns, which form sperm flagella, than other species having vascular systems but lacking flagella during evolution (Supplementary Table S2).

### Genes for thermogenesis

To elucidate the molecular basis of cone thermogenesis, we performed transcriptome analysis during the diurnal thermogenic period. We identified 89 upregulated and 36 downregulated transcripts in thermogenic tissues (Supplementary Figure S7). Central to this heat-production mechanism was the identification of Alternative Oxidase 1 (AOX1). Three of the upregulated candidates, including AOX1, were predicted to possess mitochondrial transit peptides, supporting their role in diverting redox energy from the respiratory chain to heat production. These findings provide a direct link between genomic plasticity and the specialized physiological adaptations of cycads.

## Discussion

We present the chromosome-scale assembly of the *C. revoluta* genome spanning 11.6 Gb in length, of which 11.4 Gb was anchored to chromosomes as pseudomolecules (Figure 1, Table 1). This assembly represents the second chromosome-scale genome for the order Cycadales, enabling a direct comparison with the closely related *C. panzhihuaensis*^2^ (Figure 2A). The *C. revoluta* chromosome-scale assembly (11.4 Gb) is 1.2 Gb longer than that of *C. panzhihuaensis* (10.2 Gb), a difference primarily attributable to repeat expansion at the distal ends of chromosomes, as no large-scale structural rearrangements were detected (Figure 2A). While the two species share a high percentage of orthologous gene clusters (90.6%), each possesses a distinct set of unique gene clusters (4.9% in *C. revoluta* and 4.5% in *C. panzhihuaensis*) that likely arose after their diversification approximately 19-20 million years ago (Figure 2B). This highlights a central paradox in cycad evolution: although they are an ancient group of ‘living fossils’, many cycad genera exhibit high species diversity resulting from more recent divergences. Further genomic research on cycads is needed to reveal the molecular mechanisms that enable both the conservation of a long-lived lineage and a species diversification system comparable to that of angiosperms.

Another key feature of this genome assembly was its extraordinary repeat content; repetitive sequences occupy 9.8 Gb (84.2%) of the assembly, with LTR retrotransposons alone constituting half of these repeats (Supplementary Table S1). The intraspecific genetic variation in *C. revoluta* revealed that both SNPs and SVs were biased toward intergenic regions (Figure 3A). This pattern likely reflects the prevalence of neutral mutations in intergenic regions and the impact of purifying selection against deleterious mutations in genic regions. In contrast, large indel polymorphisms (5,000 to 20,000 bp) associated with LTR retrotransposons showed a relative bias toward genic regions, including untranslated regions (UTRs) and introns. The predominance of low-frequency transposon insertion polymorphisms (Figure 3B) indicates that a recent increase in the activity of these insertions and deletions occurred during the distribution expansion, rather than being maintained at intermediate frequencies through long-term neutral evolution.

The genomic localization of TEs in plants has been documented in several pangenome studies of model organisms, such as *A. thaliana* and rice, and two non-mutually exclusive possibilities have been discussed ^65–67^. First, the initial TE insertions may occur randomly, with their subsequent retention being favored by natural selection if they contribute to gene expression regulation and individual survival. Second, TE targeting could be directed by specific chromatin signatures, such as histone modifications that characterize transcriptionally active and accessible chromatin. These epigenetic landmarks serve as recruitment signals for new TE insertions toward genic regions and TE deletions by homologous recombination. Transposon insertion polymorphisms in UTRs and introns, which have been associated with local adaptation in angiosperms, may also occur in gymnosperms, despite their substantially larger genome sizes and higher proportions of intergenic regions compared to angiosperms. These results imply that these mechanisms may be involved in adaptive evolution, although further functional studies are needed to confirm their role in speciation.

We constructed a continuous assembly of the sex chromosome, including the male-specific region (MSR) at the end of the chromosome 8. The sex-determining region in *C. revoluta* exhibited complex repetitive patterns expected to result in recombination suppression (Supplementary Figure S5). This structure suggests that the mechanism of sexual dimorphism maintenance is the XY system. Cycads and ginkgos are dioecious gymnosperms that possess a unique male trait intermediate between basal and other seed plants, specifically the production of large, multi-flagellated motile sperm within pollen tubes. Because the genes for sperm flagella and those expressed in male organs in *C. revoluta* are mainly located on autosomes, the sex-determination genes in the MSR could trigger the production of sperm flagella and male-specific traits.

Moreover, the phylogenetic analysis revealed species-specific MADS-box gene clusters in cycads and ginkgos. Tandem duplication and faster birth-and-death evolution of MADS-box genes are well known in model land plants^68,69^. These species-specific gene clusters in cycads and ginkgos might have emerged after rapid radiations ranging from 11 to 20 million years ago and might contribute to the development and structural diversification of reproductive organs. Our findings will contribute to understanding the unique sexual dimorphic traits and the evolutionary history of sex chromosomes from seedless plants to angiosperms.

We detected alternative oxidase (AOX) as a candidate associated with thermogenesis, consistent with a recent study in *Zamia furfuracea*, a cycad belonging to a different family^70^. The AOX detected in this study had different sequences and genomic positions from those of CrAOX1 and CrAOX2 reported in the previous study^9^, whereas its sequence was more similar to CrAOX1 than to CrAOX2. AOX1 is a transmembrane protein localized to the inner mitochondrial membrane, has mitochondrial transit peptides, and bypasses the proton-translocating multi-protein complexes^71,72^. Of the candidates, three genes with mitochondrial transit peptide, including those of unknown function, might also be associated with thermogenesis. This suggests that thermogenesis was comprehensively driven by multiple genes associated with the mitochondrial electron transport chain and cellular respiration^73^. The previous study detected circadian clock genes, but our study did not. The different result is reasonable because *C. revoluta* does not clearly exhibit a circadian rhythm in thermogenic activity, unlike the genus *Zamia*^9,70^.

In conclusion, this high-quality genome resource establishes *C. revoluta* as a key model organism, directly resolving the genomic architecture underlying cycad diversification and species diversity. Crucially, the genomic data also offer vital genetic tools to mitigate the devastating cycad aulacaspis scale (*Aulacaspis yasumatsui*) damage currently threatening cycads in Okinawa^74^, while providing strategies to safely manage or reduce cycasin^75^, the toxin responsible for severe historical crises known as “cycad hell.” Ultimately, protecting this lineage through genomic insights ensures the survival of both an ancient evolutionary wonder and a deeply rooted ethnoecological heritage.

## Acknowledgments

We thank the Koishikawa Botanical Garden (The University of Tokyo) for providing the plant materials. We also thank K. Ozawa, C. Minami, H. Tsuruoka, Y. Kishida, and A. Watanabe (Kazusa DNA Research Institute) for their technical assistance.

## Supplementary Data

**Supplementary Table S1.** Repetitive sequences in the *C. revoluta* genome assembly.

**Supplementary Table S2.** Counts of candidate genes of seven categories involved in sperm flagellum biosynthesis in 14 species.

**Supplementary Figure S1.** Relationship between markers obtained by pollen RNA-seq on Omni-C scaffolds.

The lower triangle of the matrix shows correlation coefficients between markers, while the upper triangle shows distances between marker positions.

**Supplementary Figure S2.** A genetic map of *C. revoluta*.

Left: SNP loci on the genetic map (vertical lines) and physical map (bars) are connected with horizontal lines. Right: Positions of SNP loci are indicated with dots on the genetic map (y-axis, cM) and physical map (x-axis, Mb). The vertical yellow lines indicate boarders of contigs.

**Supplementary Figure S3.** Distribution of telomere repeats on each pseudomolecule with 1-Mb windows.

TTTAGGG and TTCAGGG are counted as plus, and CCCTAAA and CCCTGAA are counted as minus.

**Supplementary Figure S4.** Sequencing depth on whole genomic regions of male and female individuals from Miyazaki, Kisarazu, and Tokyo.

Red lines indicate the mean depth of the whole genome.

**Supplementary Figure S5.** Dot plot of sex-determining regions. Line colors represent sequence identity.

A. Comparison of sex chromosomes and sex-determining regions between *C. panzhihuaensis* and *C. revoluta*. (B) Self-alignment of the X contigs in *C. revoluta*. Identical sequences are ignored. (C) Self-alignment of the initial 60 Mb of the sex chromosome in *C. revoluta*. Identical sequences are ignored.

**Supplementary Figure S6.** Phylogenetic relationship of MADS-box transcription factor genes in *C. revoluta*, *C. panzhihuaensis*, and *G. biloba*.

**Supplementary Figure S7.** Transcriptome analysis of thermogenesis-associated genes.

A. A Venn diagram of the number of differentially expressed genes. (B) A heatmap of differentially expressed genes associated with thermogenesis. The x-axis indicates microsporophylls (ML) and microsporangia (MG) collected at early thermogenic (Early), middle thermogenic (Middle), and late non-thermogenic (Late) stages.

## Data availability

Raw sequence reads and assembled sequences were deposited in the DNA Data Bank of Japan (DDBJ) BioProject database under the accession number PRJDB42334. The genome assembly files and gene annotation files are available in Kazusa Genome Atlas (https://genome.kazusa.or.jp).

