## Supplementary Figure for "Chromosome-scale genome assembly of *Cycas revoluta* provides insights into cycad diversification and species diversity"

1. Department of Frontier Research and Development, Kazusa DNA Research Institute, Chiba 292-0818, Japan
2. School of Marine Bioscience, Kitasato University, Kanagawa 252-0373, Japan
3. Department of Biological Science, Graduate School of Science, The University of Tokyo, Tokyo 113-0033, Japan
4. Faculty of Agriculture, University of Miyazaki, Miyazaki 889-2192, Japan
5. Advanced Genomics Center, National Institute of Genetics, Shizuoka 411-8540, Japan
6. Graduate School of Environmental and Life Science, Okayama University, Okayama, Japan
7. Graduate School of Life Science, Tohoku University, Miyagi 980-8577, Japan
†. Present address: Department of Frontier Research, National Institute of Genetics, Shizuoka 411-8540, Japan

**Supplementary Table S1.** Repetitive sequences in the *C. revoluta* genome assembly.
**Supplementary Table S2.** Counts of candidate genes of seven categories involved in sperm flagellum biosynthesis in 14 species.

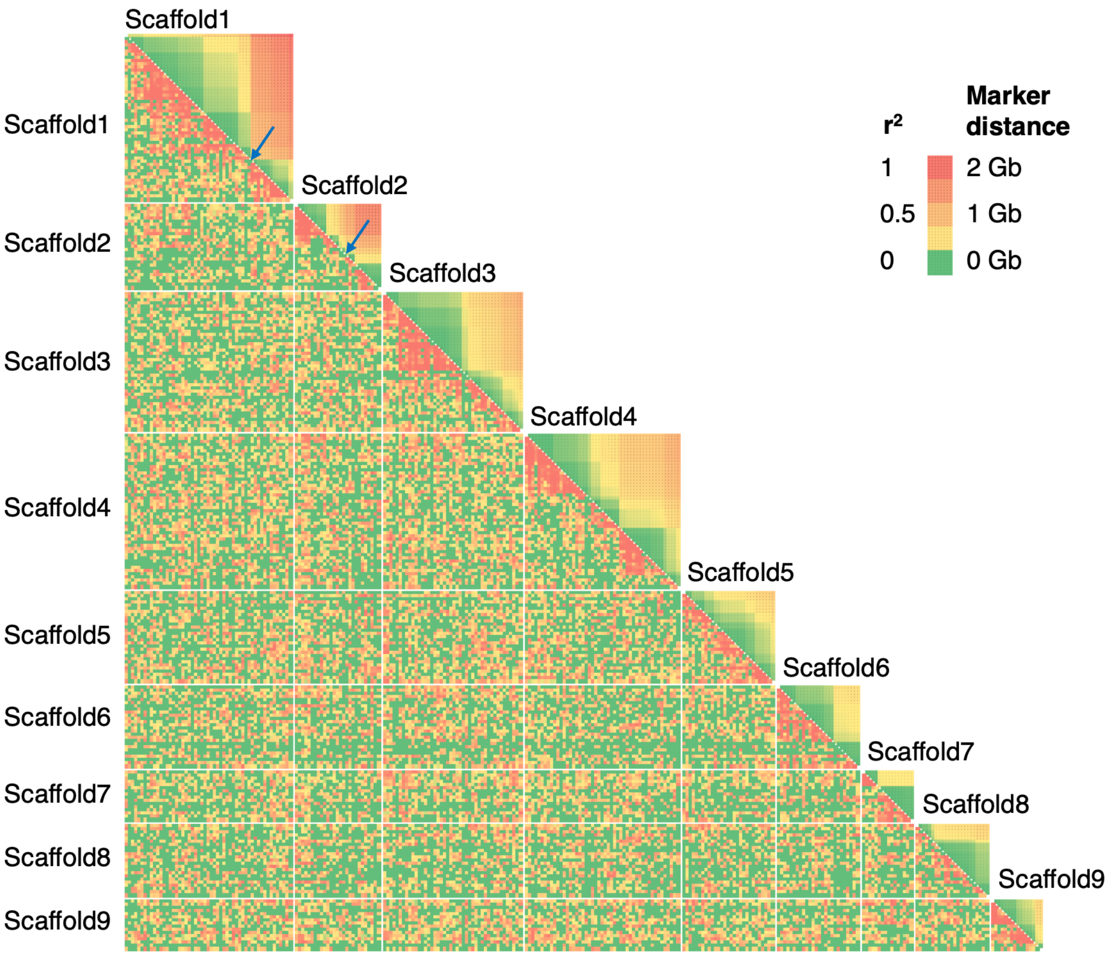


**Supplementary Figure S1.** Relationship between markers obtained by pollen RNA-seq on Omni-C scaffolds.

The lower triangle of the matrix shows correlation coefficients between markers, while the upper triangle shows distances between marker positions.

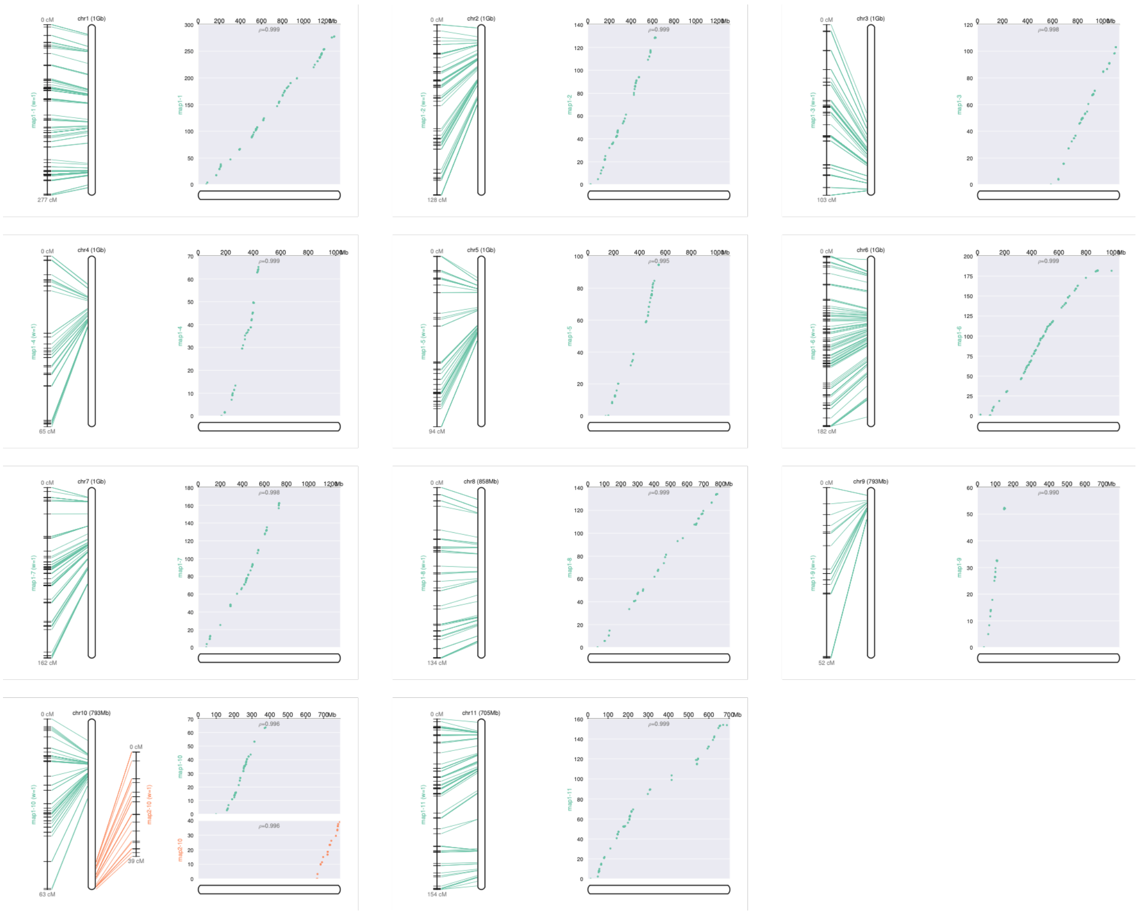

**Supplementary Figure S2.** A genetic map of *C. revoluta*.

Left: SNP loci on the genetic map (vertical lines) and physical map (bars) are connected with horizontal lines. Right: Positions of SNP loci are indicated with dots on the genetic map (y-axis, cM) and physical map (x-axis, Mb). The vertical yellow lines indicate boarders of contigs.

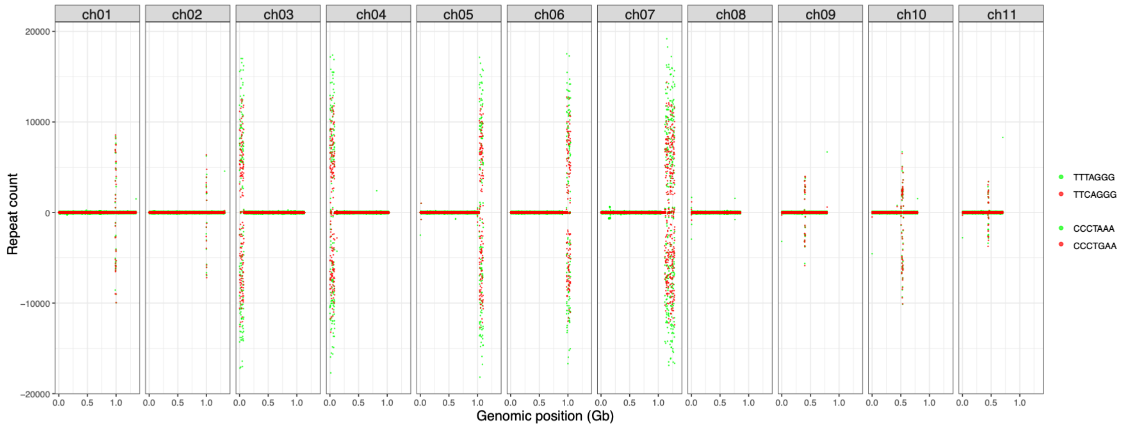


**Supplementary Figure S3.** Distribution of telomere repeats on each pseudomolecule with 1-Mb windows.

TTTAGGG and TTCAGGG are counted as plus, and CCCTAAA and CCCTGAA are counted as minus.

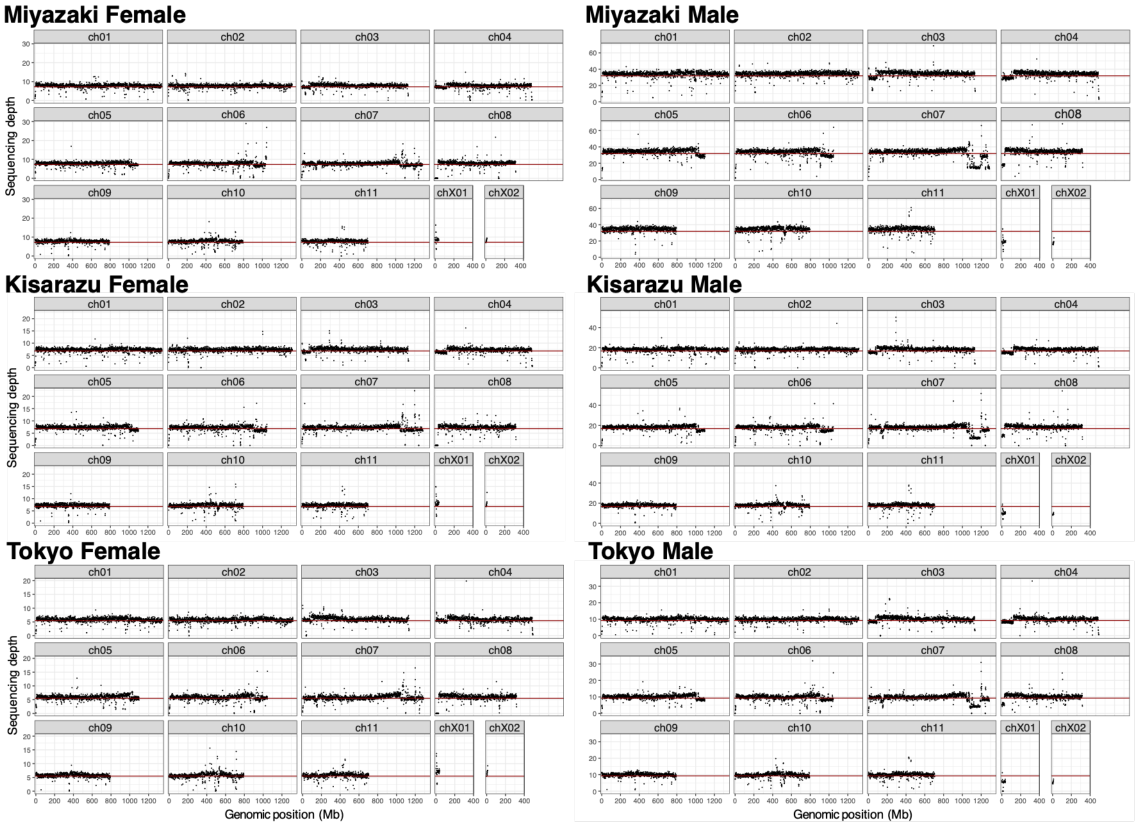


**Supplementary Figure S4.** Sequencing depth on whole genomic regions of male and female individuals from Miyazaki, Kisarazu, and Tokyo.

Red lines indicate the mean depth of the whole genome.

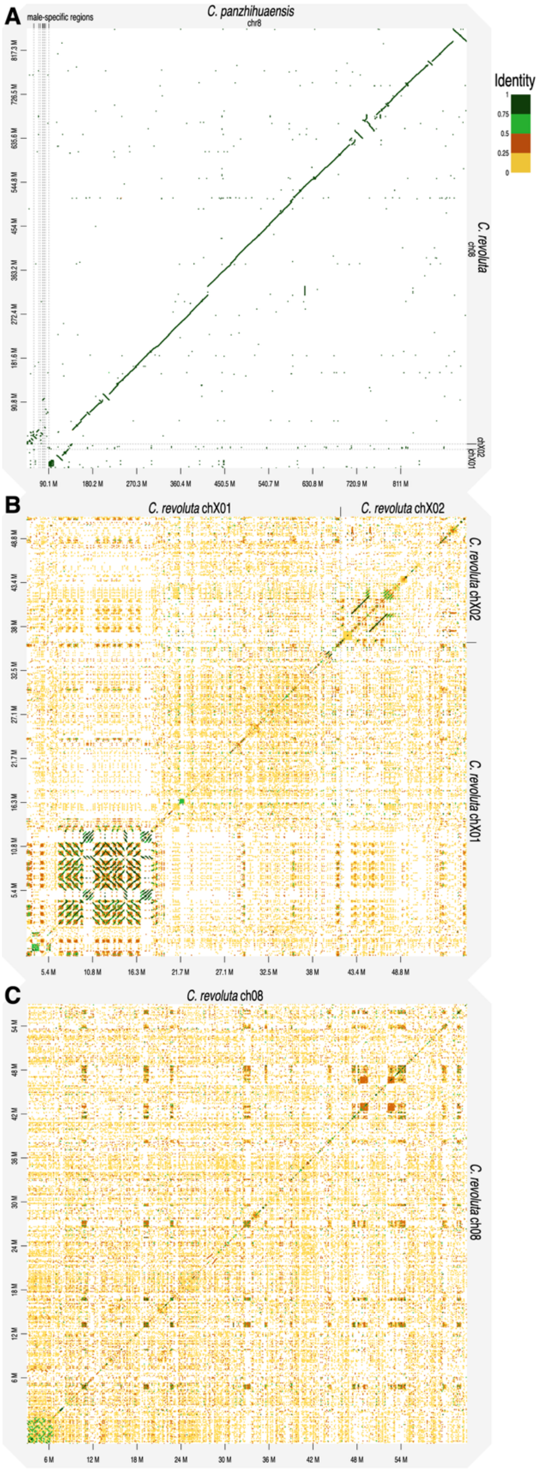


**Supplementary Figure S5.** Dot plot of sex-determining regions.

Line colors represent sequence identity. (A) Comparison of sex chromosomes and sex-determining regions between *C. panzhihuaensis* and *C. revoluta*. (B) Self-alignment of the X contigs in *C. revoluta*. Identical sequences are ignored. (C) Self-alignment of the initial 60 Mb of the sex chromosome in *C. revoluta*. Identical sequences are ignored.

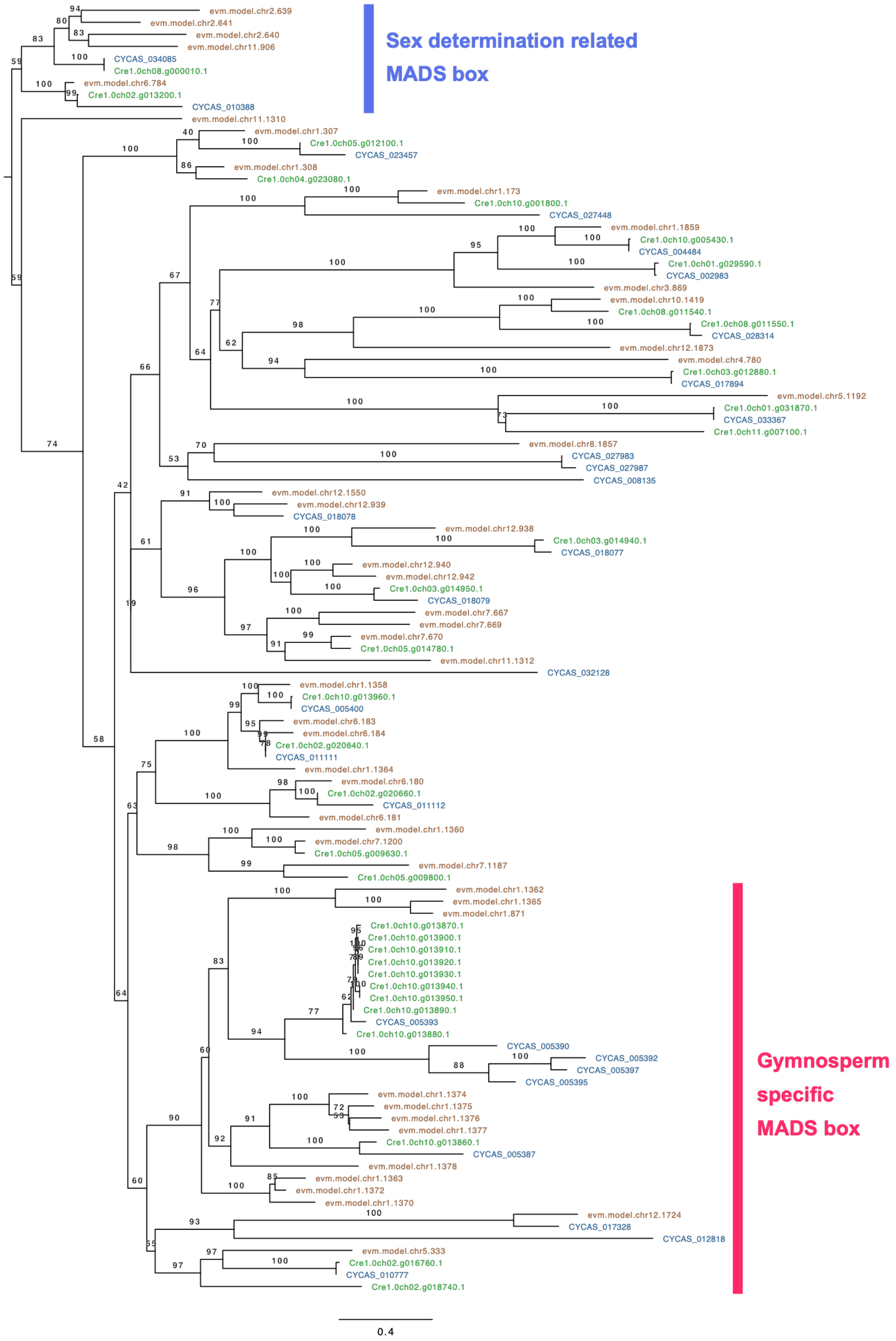


**Supplementary Figure S6.** Phylogenetic relationship of MADS-box transcription factor genes in *C. revoluta*, *C. panzhihuaensis*, and *G. biloba*.

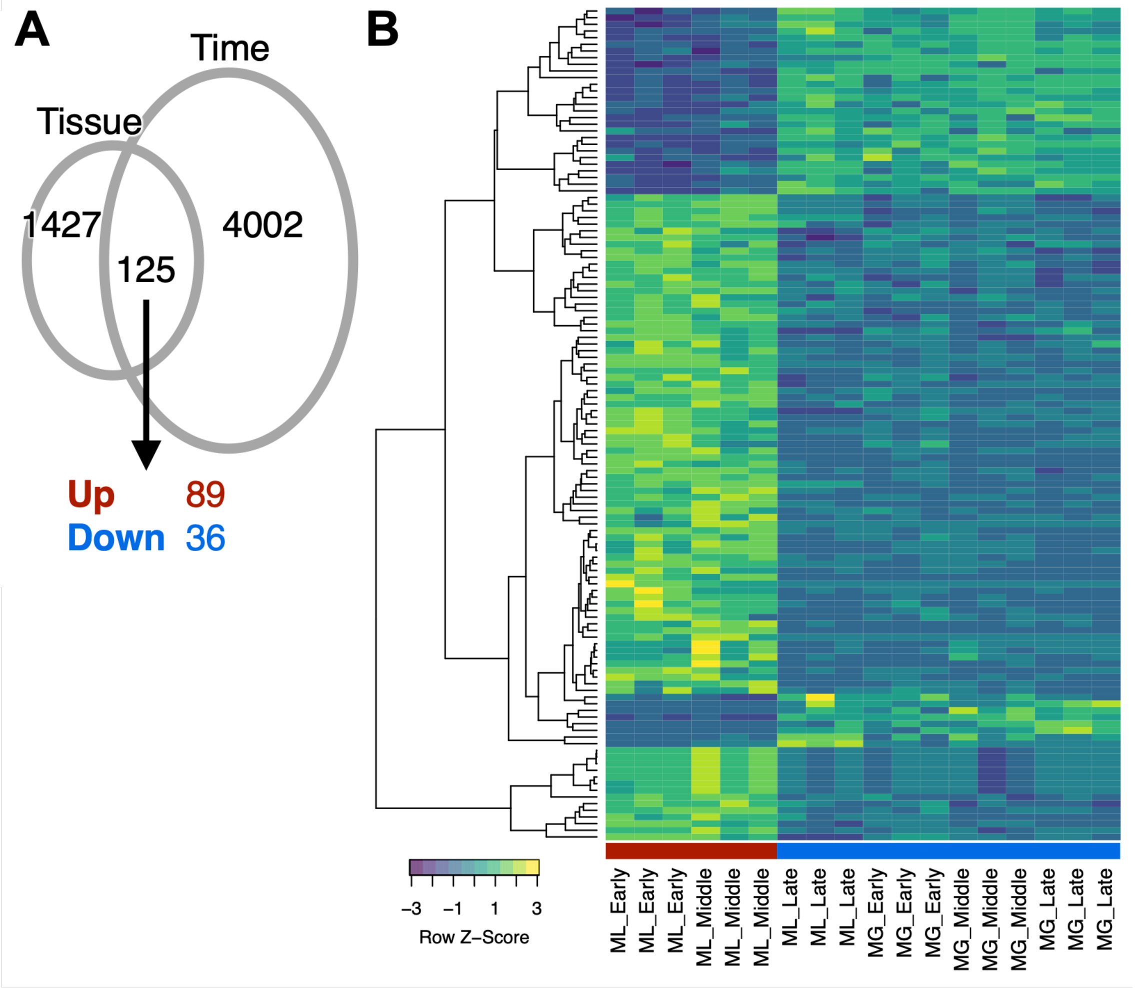


**Supplementary Figure S7.** Transcriptome analysis of thermogenesis-associated genes. (A) A Venn diagram of the number of differentially expressed genes. (B) A heatmap of differentially expressed genes associated with thermogenesis. The x-axis indicates microsporophylls (ML) and microsporangia (MG) collected at early thermogenic (Early), middle thermogenic (Middle), and late non-thermogenic (Late) stages.
